# 4D chromosome reconstruction elucidates the spatial reorganization of the mammalian X-chromosome

**DOI:** 10.1101/2021.01.21.427652

**Authors:** Anna Lappala, Chen-Yu Wang, Andrea Kriz, Hunter Michalk, Kevin Tan, Jeannie T. Lee, Karissa Y. Sanbonmatsu

## Abstract

Chromosomes are segmented into domains and compartments; yet, how these structures are spatially related in 3D is unclear. Here, by directly integrating Hi-C capture experiments and 3D modeling, we use X-inactivation as a model to examine the time evolution of 3D chromosome architecture during substantial changes in gene expression. First, we show that gene expression A/B compartments are consistent with phase separation in 3D space. Second, we show that residuals of smaller scale structures persist through transitions, despite further large-scale reorganization into the final inactive configuration, comprising two “megadomains”. Interestingly, these previously hidden residual structures were not detectable in 2D Hi-C maps or principal component analyses. Third, time-dependent reaction-diffusion simulations reveal how Xist RNA particles diffuse across the 3D X-superstructure as it reorganizes. Our 4DHiC pipeline helps satisfy the growing demand for methodologies that produce 3D chromosome reconstructions directly from 2D datasets, which are consistent with the empirical data.

## Introduction

Chromosomes exhibit complex non-equilibrium dynamics. In many cases, their structural reorganization contributes to important physiological transitions in the cell. The nature of chromosome dynamics is poorly understood and requires a detailed representation of its 3D structural ensemble as it evolves over time. Such 3D data is also critical for understanding the extent to which gene expression and epigenetic mechanisms are regulated by 3D architecture in the context of chromatin ^1-2^. Breakthrough technological advances such as “Hi-C” have enabled large-scale sequencing of genome-wide 2D contacts ^3^ and validated earlier observations that mammalian chromosomes are organized into loop domains, anchored by CCCTC-binding factor (CTCF) proteins, and into larger compartments, which roughly correlate with gene expression states ^4-9^. While it is well-established that long-range loops between regulatory elements such as enhancers and promoters play a role in developmentally timed gene expression ^10-12^, studies have also shown that abolition of topological domains and major shifts in compartmentalization have little immediate impact on gene expression ^13-19^. These studies have therefore left open the question of whether and how chromosome structure regulates gene expression. New methodologies are required to push the present boundaries of knowledge. While there are models that demonstrate important specific features of chromosome architecture ^20-22^, critically missing are methodologies that *directly* convert 2D biological information (such as from Hi-C contact maps) into 3D models of the chromosome, thus providing a generalized experiment-driven methodology to model chromosome structure. Such methodology would yield direct visualization of empirically measured architectural information in 3D. Here, we develop new enabling models for 3D reconstruction and analysis of time evolution, the result of which is a 4D map of the classic epigenetically regulated chromosome — the inactive X (Xi) of the female mammal.

A number of 3D models have been developed over the past decade ^23 22 24^, and these can be roughly classified into two groups: ‘mechanistic’ and ‘inverse’ models (for a full review see ^25^). Mechanistic models are used to test hypotheses for underlying mechanisms such as microscopic DNA properties, mechanisms of chromosome compartment- and domain formation, loops and loop domain formation and dynamics. ^26,27^ These models also provide detailed structures of gene loci. ‘Inverse’ models, on the other hand, use data to infer chromatin structures, generating *in silico* configurations of gene loci or whole chromosomes consistent with HiC data. Our models, which use our 4DHiC pipeline (described below) fall into the ‘inverse’ model category. A prominent example of a constraint-based model includes TADbit, which has been used to reconstruct gene loci in mammalian cells^24^,. Similar methods have been applied to reconstruct whole yeast nuclei.^28^ In polymer-based models, iterative parameter determination has been combined with both Monte Carlo and Molecular Dynamics (MD) simulations to generate structures from 5C and Hi-C data. By reducing model resolution, whole human chromosomes or even whole nuclei can be modelled and configurations from single-cell Hi-C data have been reported. ^29^ A key advantage of 4DHiC is that it makes no assumptions regarding the structure of the chromosome. For example, we do not assume A compartments interact with A compartments, nor that certain genes or regions are critical in structure formation, nor that CTCF is important for loop formation. In addition, we do not rely on the approximation that the distance between Hi-C interacting beads is inversely proportional to their interaction frequency, which can create distortions in the chromosome structure by forcing beads with weak interaction strengths to be far apart when in actuality, they may be just beyond cross linking distance but not far apart. Instead, we use an automated procedure to read in the experimentally determined Hi-C map and produce a corresponding 3D model. The features of our 3D reconstructions emerge from the experimentally determined Hi-C data itself and do not require calling of TADs, important genes or other human-readable features of Hi-C maps. By using harmonic constraints to simulate cross linking distances in the context of polymer simulation, we achieve direct, experimentally validated correspondence between the 3D model and the experimentally determined Hi-C contact map, allowing us to visualize complicated features observed on 2D Hi-C map in 3D without prior assumptions about any features in the Hi-C maps. The 4DHiC pipeline is transferable in the sense that it can be applied by a user to any Hi-C map.

In female mammals, one X-chromosome is transcriptionally inactivated to balance gene expression between females and males. X-chromosome inactivation (XCI) involves a progressive transformation of an active X chromosome (Xa) to an inactive X chromosome (Xi), precipitated by Xist RNA molecules spreading over the chromosome ^30-33^. Many 1D (*e*.*g*., epigenetic sequencing) and 2D (*e*.*g*., Hi-C) studies of X inactivation have been performed and together they point to three evolving principles of chromosome organization (Fig. 1A): (i) A/B compartments, delineating active and inactive regions ^8^, (ii) S1/S2 compartments, appearing during the transition from Xa to Xi ^34^, and (iii) two megadomains, a structural endpoint for the Xi ^8,19,35-38^. These studies have complemented a wealth of microscopy studies regarding the large-scale organization of the Xi in the nucleus ^39-45^. While these earlier studies have provided valuable ideas, many open questions remain. What is the relationship between A/B and S1/S2 compartments? Are S1/S2 structures transitory or do they persist during XCI? How and why do megadomains form? Finally, how does Xist RNA spread over the evolving 3D Xi geometry? Here, we address these questions by employing our 4DHiC pipeline, producing experimentally validated 3D models from experimentally determined Hi-C maps describing various stages of X inactivation. To understand the spreading of Xist RNA molecules around and through the X chromosome during inactivation, we perform particle-based reaction diffusion simulations.

**Fig 1.**
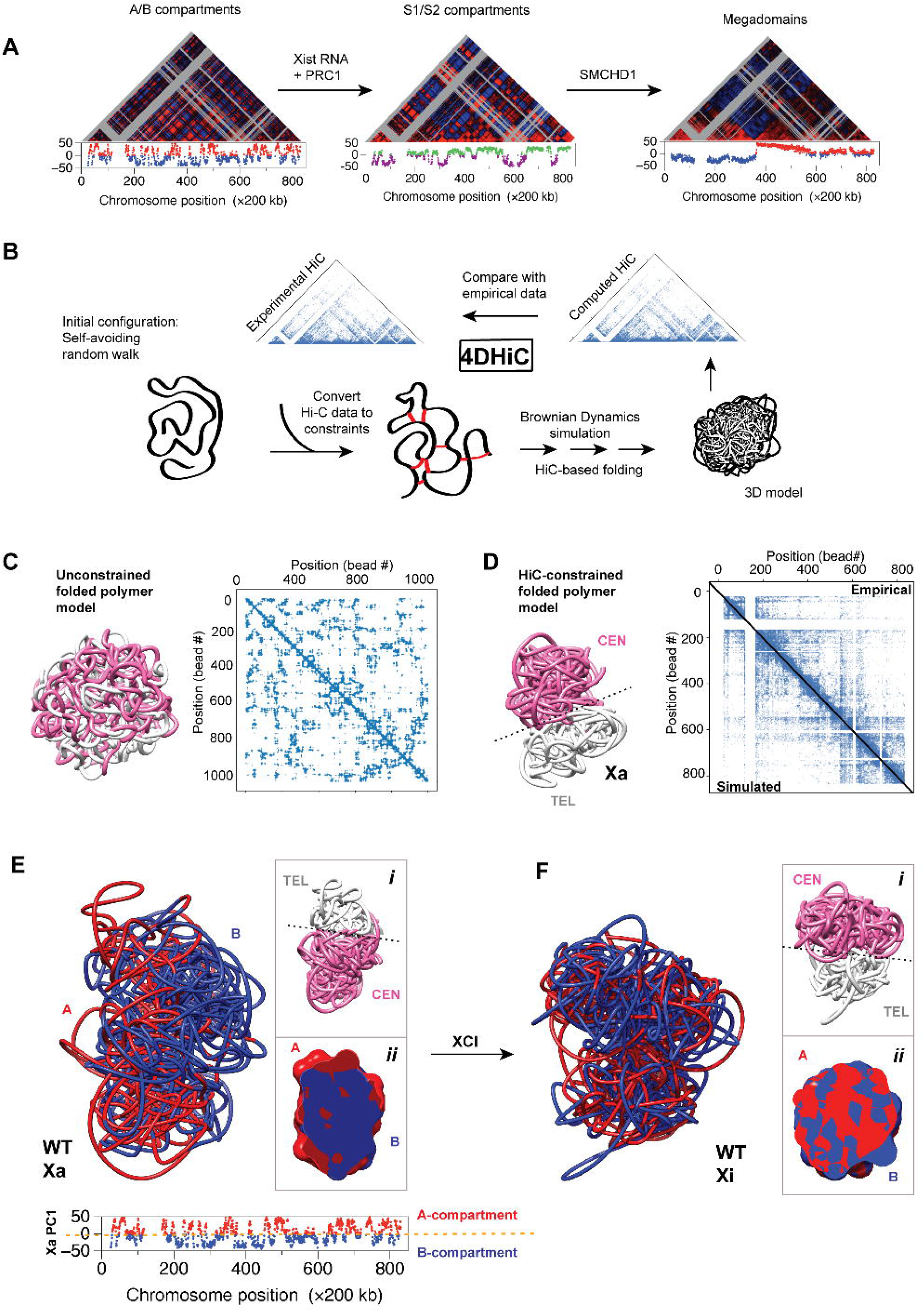
3D reconstructions of the active (Xa) and inactive (Xi) X-chromosomes, based on 2D Hi-C experiments, show that megadomain formation entails spatial dismantling of A/B compartments. **(A)** Schematic of megadomain formation during X-chromosome inactivation (XCI) showing Pearson correlation maps and the first principal components (PC1) at each stage. (Adapted from ^34^) **(B)** Schematic of 4DHiC pipeline: virtual crosslinks are created from experimentally determined Hi-C maps and applied to beads-on-string model of chromosome with random 3D structure; MD simulations with constraints produce a folded chromosome structure; simulated Hi-C map is computed from folded structure; simulated map is compared to experimentally determined map; MD simulations proceed until convergence between simulation and experiment is achieved. **(C)** Results of the unconstrained folded polymer model (folded equilibrium globule) shows mixing of ‘centromeric’ and ‘telomeric’ regions (pink and white, respectively). Contact map for a folded equilibrium globule shows extensive mixing within the chain. The matrix is divided into two triangles, with each showing the same pairwise comparison and data taken from an unconstrained “randomized” model. **(D)** 3D model of the wild type Xa. White: region of sequence corresponding to telomeric megadomain. Pink: region of sequence corresponding to centromeric megadomain. Comparison between experimental (upper triangle) vs. simulated (lower triangle) Hi-C contact maps for the WT Xa chromosome shows agreement. **(E)** A/B compartments defined by principal component (PC) analysis of Xa projected onto 3D reconstruction of Xa shows spatial separation between A and B compartments in physical 3D space. Red: positive values of first principal component (PC1); blue: negative values of PC1. Inset (i): regions corresponding to megadomains (colors same as (D). Inset (ii): cross section. Lower inset: PC1 vs. 1D sequence position shows that A and B compartments are mixed in 1D sequence space. **(F)** 3D reconstruction of Xi showing mixing of A and B compartments in 3D physical space (colors same as (E)) Inset (i): regions corresponding to megadomains (colors same as (D). Inset (ii): cross section.

## Results

### 3D Reconstructions from HiC with 4DHiC

We generate 3D reconstructions of the X chromosome directly from experimental 2D Hi-C contact maps (Movie 1). Because there are many possible 3D folds consistent with the 2D experimental data, there are a variety of approaches to solve this problem. In other words, connectivity information alone does not provide all the information needed to generate a 3D structure because only relative positions (in terms of contacts) are defined, as opposed to directly ‘measured’ 3D structural information. Previous efforts have employed polymer simulations from first principles with great success, incorporating well-defined 1D information such as CTCF motifs, reconstructing 2D connectivity profiles from first principles ^21,22,46^. This approach has been successful in identifying critical elements of chromatin organization such as loops, leading to the proposed mechanism of chromatin folding by loop extrusion ^22^. These previous models performed by other groups provide a possibility to generate a contact map without the need to perform Hi-C experiments in the first place, assuming that the 1D information is sufficient for such predictions, which could be the case due to universality of certain motif interactions, as in the case of CTCF. However, there could be yet unknown factors and motifs that might change the model as new experimental techniques emerge. Another type of proposed model is the so-called minimal chromatin model ^24^ whereby the genome is partitioned into groups of a several types, such that each type of group is marked by characteristic histone modifications and interacts with a characteristic combination of nuclear proteins. These models require a number of experimentally motivated assumptions and a high number of fitting parameters, which may thereby affect the transferability of models of such kind. For example, stipulating that regions of chromatin with similar epigenetic profiles would interact is a reasonable assumption. However, other possibilities could occur, such as simultaneous silencing of regions distant from each other in 3D space, cannot be excluded. To minimize assumptions and biases, here we ask the questions: Given Hi-C data, can a model be generated such that all 2D biological information is mapped directly onto the 3D structure without assumptions and prior knowledge of the epigenetic modification state of the chromosome, and can a modeling pipeline be developed that would be transferable to any Hi-C dataset?

Here, we present a pipeline (4DHiC) to produce 4D chromosome reconstructions (i.e., temporal series of 3D chromosome configurations) directly from 2D datasets, which are consistent with the empirical data. Our approach uses a coarse grained polymer model. The approach is entirely scalable: the number of beads can be increased or decreased, allowing tuning of the resolution of the polymer to the resolution of any Hi-C dataset (Fig. 1B). The approach does not rely on the multidimensional scaling approximation (MDS), which infers 3D distances based on measured contact frequency.^47^ We adapted the polymer bead-spring model from Kremer and Grest – a widely used polymer physics model that describes a polymer as beads connected by springs^48,49^. Our Hi-C datasets for X inactivation have a resolution of 200 kb. Thus, we model the 166-Mb X chromosome as a polymer with 833 beads, corresponding to 200 kb resolution, with one bead for each Hi-C bin. The initial self-avoiding random walk (SARW)-based polymer model is collapsed by constraining strongly attracting monomers, imitating pairwise cis-interactions captured by Hi-C experiments. Simulations are continued until convergence is achieved between simulated 2D contact maps and experimentally determined 2D Hi-C maps (correlation coefficient *R* ∼ *0*.*97*) (Fig. 1).

To validate our 4DHiC pipeline, we first reconstructed the active X (Xa) and colored the chromosome into two halves separated by the microsatellite repeat, *Dxz4* ^8,17,19,37,38,50^— a centromeric (CEN) half in white corresponding to the proximal megadomain on the Xi (0-72 Mb) and a telomeric (TEL) half corresponding to the distal megadomain on the Xi (73-166 Mb) in purple (Fig. 1). Without constraints, the model polymer chain folded into a 3D structure without apparent organization (Fig. S1), as expected, with the two halves (white, purple) of the Xa fully mixed in 3D (Fig. 1C, left). On the other hand, constraining the model using empirically measured 2D Hi-C contacts generated an orderly 3D chromosome model, with the proximal and distal halves of the Xa separated into two poles (Fig. 1D, left). The simulation-derived 2D contact map correlates strongly with the Hi-C-derived map (*R*_*cp*_ *= 0*.*97, R*_Pearson_ = 0.85, Fig. 1D, right). In particular, although the long-range interactions required for Xi megadomains are not present in the Xa, the two chromosomal halves still undertook compact globular morphologies that were spatially separated from each other in 3D space, consistent with previous studies ^41^. Altogether, these data validate our pipeline.

We then asked whether the 4DHiC pipeline could model the multi-megabase, alternating “A” and “B” compartments that segregate mammalian chromosomes on the basis of chromatin states which can be detected by principal component analysis (PCA) of Hi-C data, often correlated with gene expression states. In the first principal component (PC1) of the Xa, active chromatin roughly correlates with the “A” compartment (eigenvectors assigned to positive values), while silent chromatin roughly correlates with the “B” compartment (negative values) (Fig. 1E, bottom inset) when the PCA was performed with 200-kb resolution. For each region of the chromosome (*i*.*e*., bin of the Hi-C map), a connectivity profile is determined (row of contact matrix); a covariance matrix is calculated from covariation of profiles; in the initial basis set, each dimension corresponds to a chromosome region (bin); the covariance matrix is then diagonalized by finding a new basis set (consisting of linear combinations of regions) that maximizes the collective variation of profiles with across the chromosome. Essentially, the first eigen vector (PC1) is composed of a linear combination of a special subset of bins whose connectivity profiles are the most highly coupled. This subset includes, but it not limited to, boundaries between spatially localized domains. The collective coordinate (PC1) often couples many regions that a distant from each in sequence space but span the entire chromosome. Thus, although chromosomes are segmented into dozens of A/B compartment regions, PC1 data are 1D. Thus, how compartments are spatially related in 3D is not immediately clear from the PC1 track. The ability to visualize this spatial connectivity would be beneficial for structure-function analysis.

Our first challenge was to map the A/B compartments derived from PC1 of Xa (Fig.1E, bottom) to a 3D reconstructed model of the Xa. Here, positive PC1 values represent A compartments, which are strongly coupled in terms of how their connectivity profiles vary with respect to one another; negative values represent the more weakly coupled B compartments. Using allele-specific HiC reads mapping to the Xa, we colored A and B regions red and blue, respectively, and mapped their positions onto the centromeric (pink) and telomeric (white) halves of our deduced Xa model (Fig. 1E). The A/B compartments of the Xa appeared somewhat segregated in 3D space into red and blue regions, with relatively little mixing between regions. Thus, the A and B compartments do not merely have opposite gene expression states and epigenetic marks but are also spatially segregated on opposite faces of the Xa in an orientation that is roughly orthogonal to the telomere-centromere axis (compare red/blue model with white/pink model; Fig. 1E, inset i). Further information could be gathered by examining tomographic sections of the Xa model. Intriguingly, there is a clear phase partitioning of A and B extending into the core of the Xa (Fig. 1E, inset ii). These models enabled visualization of the Xa compartments in 3D for the first time.

### Xi Transition from A/B to S1/S2 organization in 3D

An unresolved question is what happens to the A/B compartments during the time-dependent process of XCI. Previous work showed that, as ∼63 A/B compartment regions are reorganized on the X chromosome, ∼25 S1/S2 regions appear, directed to form by Xist RNA together with Polycomb repressive complex 1 (PRC1)^51^. The spatial relationship between A/B and S1/S2 compartments is not currently known. Are the A/B structures destroyed in order to form S1/S2 compartments? Does the chromatin relocate in order to reorganize into S1/S2 structures? From an architectural standpoint, these two scenarios would be unfavorable because of the need to destroy the pre-existing structures and to reconstruct, *de novo*, new types of structures. An alternative, a more energetically favorable mechanism might entail residuals of the smaller-scale structures persisting through the various transitions, serving as a foundation for the final Xi superstructure.

To address these questions, we first determined the structure of the Xi using allele-specific data from Hi-C experiments taken from mouse neural progenitor cells (NPCs). The Xi also partitioned in an orderly way into a centromeric and a telomeric half (Fig. 1F, inset i). However, when we mapped the A/B compartments defined from Xa onto the 3D reconstruction of Xi, there was a clear mixing of ‘red’ and ‘blue’ A/B compartments (Fig. 1F). The mixing was especially evident in a tomographic section (Fig. 1F, inset ii), in which heterogeneity contrasted with the clustered pattern seen in the A/B separation of the Xa (Fig. 1E, inset ii). These data demonstrate an apparent loss of the original A/B organization in the final Xi superstructure. This loss could be due to a large scale reorganization. Alternatively, the loss could instead suggest that the original A/B structures exist but are no longer discernible. These two possibilities cannot be distinguished by 2D contact matrices and 1D PC1 analyses, typically derived from Hi-C datasets.

To quantify mixing of A/B compartments occurring in the transition from the Xa state to the Xi state during X-inactivation, we computed a mixing ratio R_A/B_, by calculating the number of interactions between A and B compartment beads, divided by the total number of interactions made by beads in the B compartment, both with other A compartment beads and with B compartment beads. This ratio increases during X-inactivation by ∼10%, suggesting that mixing between A and B compartments occurs during inactivation (Fig. S3). As a control, a similar ratio is computed for interactions between B compartment beads with other B compartment beads. As expected, the number of these interactions decreases due to mixing with the A compartment. In effect, during X-inactivation, interactions between B compartment beads and other B compartment beads are exchanged for interactions between B compartment beads and A compartment beads.

Although the spatial organization of A/B compartments has been implied by other studies, none of them examined compartments on the Xi. It is now clear from various studies that the Xi adopts a unique conformation and therefore what is learned about autosomes cannot necessarily be extrapolated to the Xi. This is true not only of the A-A and B-B clustering, but especially true of the S1-S1 and S2-S2 co-partitioning. Our current study fills the gap in the 3D reconstruction of the Xi.

It is known that the Xi transits through at least one intermediate state prior to assuming the final Xi superstructure. To address the question of what happens to A/B chromatin, we must examine the spatial relationship between the A/B versus these transitory S1/S2 structures. The merging of S1/S2 compartments to create the Xi superstructure then depends on a non-canonical SMC protein known as SMCHD1 ^34^. While S1/S2 compartments emerge transiently on a Hi-C contact map during intermediate stages of XCI in wildtype cells, they can be seen quite clearly on an established Xi when SMCHD1 becomes deficient ^34,51-53^. That is, knocking out SMCHD1 appears to effectively stall the system in an intermediate state between Xa and Xi, exposing clear S1/S2 structures in 2-D Hi-C contact maps. The resurrected S1/S2 regions on the SMCHD1-deficient Xi are similar to those that arise as natural intermediates during XCI in wild-type cells ^34^.

We therefore turned to SMCHD1-deficient female cells with an established Xi. We produced 3D reconstructions of the SMCHD1 KO Xi and mapped the S1/S2 regions onto this mutant Xi (Fig. 2A,B). Intriguingly, the S1/S2 regions partitioned into two opposite faces on the Xi, with the S1 (green) compartment forming a continuous surface on one side of the Xi and the S2 (purple) compartment forming a continuous surface on the opposite side (Fig. 2A). Cross-sections of the 3D chromosome revealed that this organization extended beyond the surface, with S1 chromatin in one hemisphere and S2 chromatin in the opposite hemisphere (Fig. 2A, inset i). The partition occurred nearly orthogonally to the centromere-telomere axis (Fig. 2A, inset ii) and, intriguingly, also orthogonally to the axis of the two megadomains (Fig. 2A, inset ii) which persisted in spite of the SMCHD1 deficiency (Fig. 2B). The significance and the reason for nearly orthogonal partitioning in this case is not clear, and to avoid confusion and possible misinterpretation we report the result as it stands. To be more quantitative, by orthogonality we mean two vectors, say ***i*** and ***j*** in an inner product space *A* are orthogonal if and only if their inner product <***i***,***j***>=0. Geometrically, this translates to vectors being perpendicular. Notably, the simulated contact map closely approximated the empirically measured heat map (Fig. 2B, *R*_*cp*_ *= 0*.*97, R*_*Pearson*_ *= 0*.*85*). Together, these data demonstrate that the S1/S2 compartments are discretely partitioned with their axis almost orthogonal to the centromere-telomere axis and the axis of the two megadomains.

**Fig 2.**
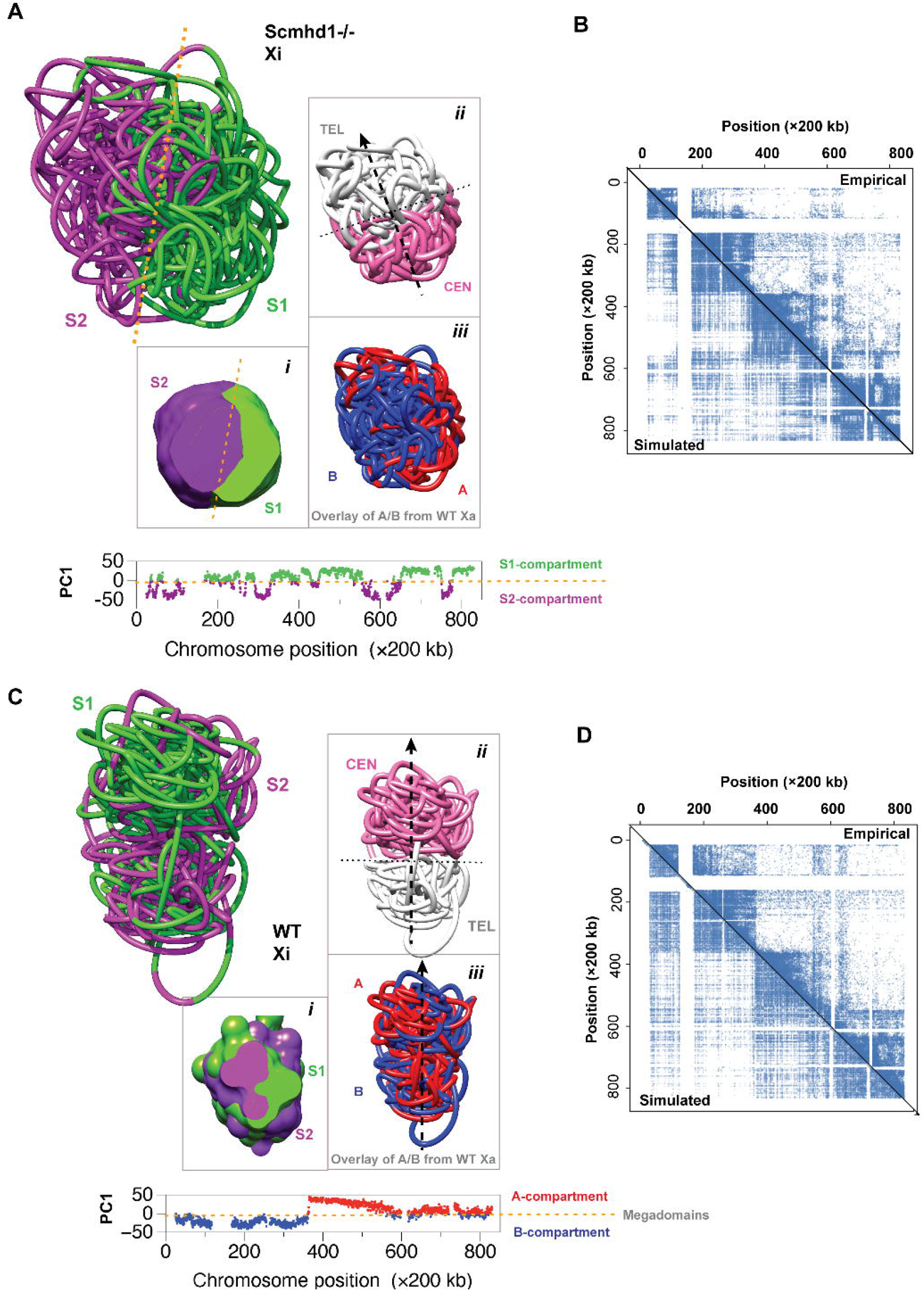
The segregation in 3D physical space of S1 (positive PC1) and S2 (negative PC1) compartments of the inactive SMCHD1 knockout chromosome based on PC analysis. **(A)** A 3D reconstruction of the SMCHD1 Xi chromosome: while S1 and S2 regions appear mixed in sequence position along the chromosome in 1D PC1 plots, 3D reconstructions reveal clear separation in 3D space. Green, positive values of first principal component (PC1); purple, negative values of PC1. Lower left inset (i): cross section; upper right inset (ii): regions corresponding to megadomains; lower right inset (iii): PC1 from Xa projected onto Xi (red, positive PC1-Xa; blue, negative PC1-Xa); bottom inset: PC1 vs. 1D sequence position shows that A and B compartments are mixed in 1D sequence space. Orange dashed lines roughly delineate separation between S1 and S2 regions. Black thick dashed line, centromere-telomere axis. Black thin dashed line, perpendicular to centromere-telomere axis. **(B)** Experimental (upper triangle) vs. simulated (lower triangle) contact map for the SMCHD1 knockout Xi chromosome. **(C)** Hi-C-based 3D reconstruction of WT inactive X chromosome: S1/S2 compartments mapped onto the WT Xi model showing mixing of S1/S2 compartments. Black thick dashed line, centromere-telomere axis. Black thin dashed line, perpendicular to centromere-telomere axis. **(D)** Experimental (upper triangle) vs. simulated (lower triangle) contact map for the wild type Xi chromosome.

We then investigated the spatial relationship between the A/B compartments found on the Xa versus the S1/S2 compartments that it acquires during XCI. We overlaid the A/B (‘red’/ ‘blue’) chromatin of the Xa onto the 3D structure of the SMCHD1 KO Xi (Fig. 2A, inset iii) and compared the red/blue (A/B) distribution to the green/purple (S1/S2) distribution. Surprisingly, the resulting overlay showed that portions of the original A/B structures remained intact beneath the S1/S2 organization. Notably, the A/B and S1/S2 organization are not identical, as indeed the ‘red’/ ‘blue’ segments cross over the axis of the S1/S2 structure. Thus, contrary to the prevailing belief that A/B organization is destroyed during XCI, it appears that elements of these original structures persist but remain hidden when viewed using conventional methodologies.

Similar questions remain unanswered for the transition from S1/S2 structures to the “compartment-less” megadomains of the Xi endpoint. Are the S1/S2 compartments destroyed in order to form the Xi superstructure? To examine the relationship between S1/S2 structures and the compartment-less Xi, we mapped S1/S2 regions of the SMCHD-deficient Xi onto the 3D model of the wild-type (WT) Xi (Fig. 2C). In contrast to the S1/S2 spatial partitioning observed for the mutant Xi (Fig. 2A), the S1/S2 (green/purple) segments as mapped onto the WT Xi showed less distinct compartmentalization in 3D physical space (Fig. 2C). Indeed, a cross section of the 3D structure showed a greater degree of S1/S2 spatial mixing when comparing the SMCHD-deficient Xi to the WT Xi (compare Fig. 2A inset i to Fig. 2C inset i). Even so, large 3D clusters of S2 (purple) and S1 (green) chromatin could still be observed. These observations suggest that S1/S2 3D organization may be less prominent, but not necessarily destroyed when the megadomain structures (Fig. 2D) form on the Xi. The megadomains identified by our simulation closely matched those empirically derived (Fig. 2D). Thus, neither the A/B nor S1/S2 3D organization appears to be entirely destroyed during the progressive Xa to Xi transformation. Echoes of these structures persist in the final Xi state and kernels of the compartments persist, possibly utilized as structural motifs in the underlying organization of the final Xi. Notably, these hidden structures were not visible in a Hi-C contact map or by principal component analysis underneath the obvious megadomains. Thus, our simulations enabled visualization of hidden spatial 3D structures that were not discernible in a conventional 2D connectivity map, thereby highlighting the power of our experiment-based structural modeling.

### Spatio-temporal evolution from Xa to Xi

While the visual aspects of 3D structures provide novel ways to interpret 1D and 2D data, importantly, they also enable the analysis of chromosome structure and gene localization dynamics based on our deduced 3D models. We first tracked changes in volume and surface area as function of time from days 0 (pre-XCI), day 3 (early XCI), to day 7 (mid-XCI) (Fig. 3A, D). During this transition, the X chromosome underwent a substantial collapse to a more compact size, both in volume and surface area, supporting the general correlation between compaction and gene silencing (Fig. S2). Interestingly, however, the surface area to volume ratio did not change dramatically during this timeframe (Fig. 3).

**Fig 3.**
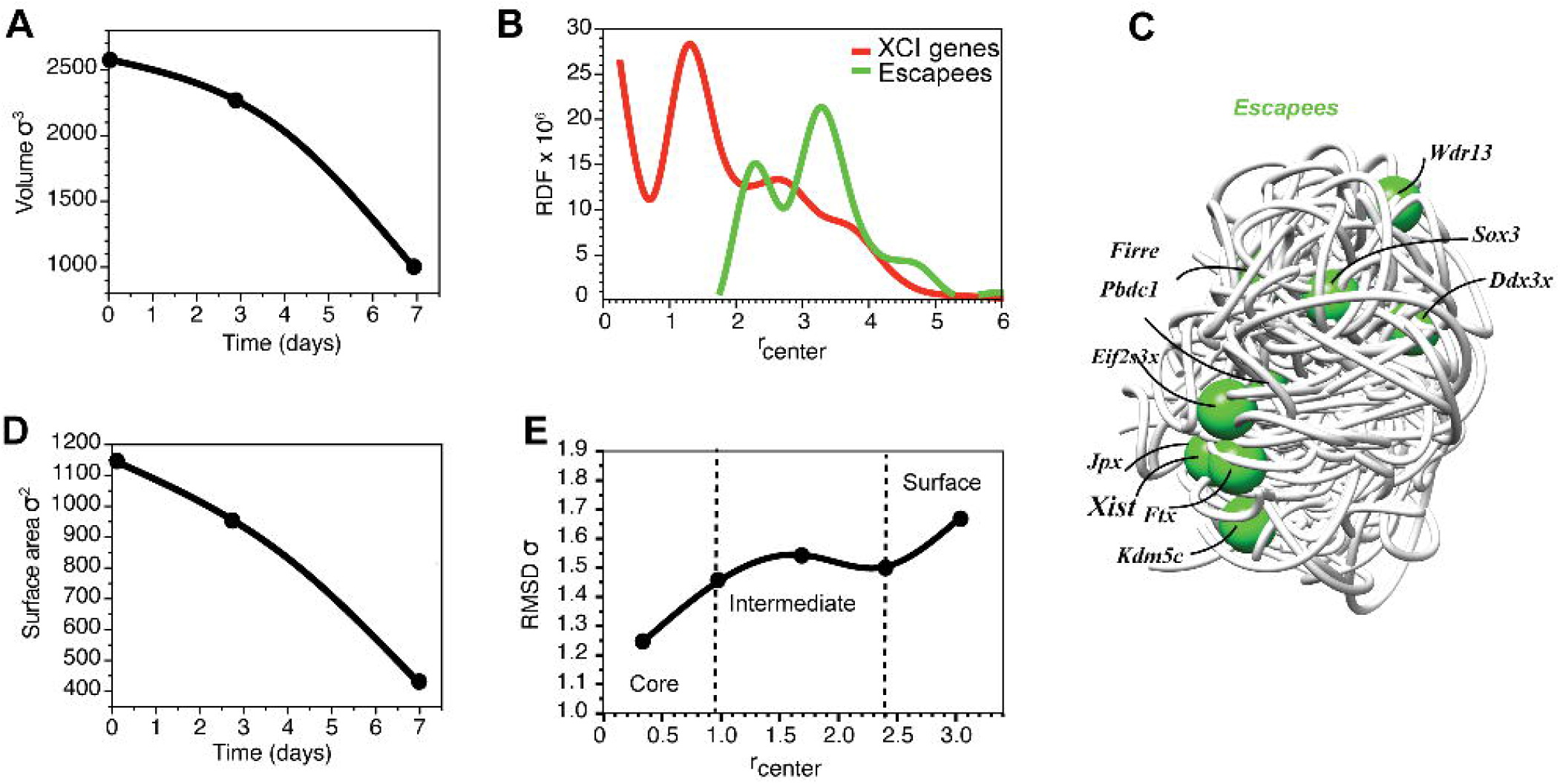
3D structural information provides critical measurements and interpretation of gene localization as well as contact probabilities. **(A)** Volume evolution over time. **(B)** Radial Distribution Function of XCI and escape genes as a function of position from the center of the WTXi chromosome taken from MEFs **(C)** 3D reconstruction of the WT inactive (WTXi) chromosome with escape genes represented as green spheres, mostly occupying the surface of the chromosome. **(D)** Surface area evolution over time **(E)** Root Mean Squared Displacement as a function of position from the center of the chromosome as the chromosome undergoes the transition from the active to the inactive state. This result demonstrates three regions of the chromosome: the dynamically arrested core, where reorganization is less likely; the mobile surface where reorganization is more likely, and an intermediate region found between the core and surface.

Next, we tracked the radial distributions of genetic elements to examine potential differential spatial distributions between genes that are subject to XCI versus the small minority of genes that escape from silencing. This radial analysis would effectively ‘dissect’ the chromosome and record whether a silent or active gene locus preferentially occupies a core, peripheral, or the surface position within the chromosome. We found that most silent genes localize in the core of the chromosome, while escapees reside in the outer regions near the chromosome surface, but not necessarily right on the surface boundary, consistent with previously published cytological data ^54 55 56^. Intriguingly, some escapees appeared to be clustered (Fig. 3C), a point not previously appreciated. Notably, recent work from many labs suggest that liquid-liquid phase separation (LLPS) and transcription hubs are critical to transcriptional activation ^57 58^. The clustering of escapee genes may occur in part due to this reason.

To compare the local radial density distributions of Xi and Xa, we calculate the corresponding radial distribution functions. Interestingly, the core was densely populated relative to the periphery (Fig. 3), suggesting that the Xi becomes denser in its core relative to the Xa. To quantitatively assess the rearrangement in 3D architecture from Xa to Xi, we calculate the root mean squared distance between particles in Xi and Xa in a shell at each radius, r_center_, and plot as a function of r_center_ (Fig. 3E). Thus, combining the analysis of radial densities and chain segment displacements during XCI progression, we observe that the chromosome is densely packed in its core, possibly too dense to be able to fully rearrange segments buried inside the core (Fig. 3E). This resembles the principle of dynamical arrest in polymer glasses (Fig. 4), and differs dramatically from the unconstrained equilibrium globule collapse (Figs. 4, S2). In order to confirm the presence of dynamical arrest in chromosomes, we looked at the Intermediate Scattering Function (the Fourier Transform of the van Hove Correlation Function) of MD trajectories of chromosome motions (Fig. 4E), which should decay to zero at long times for a liquid (no dynamical arrest), confirming that it is indeed the case for the equilibrium folded globule that behaves as a liquid droplet, however the ISF does not decay to zero for chromosomes both in their active and inactive states, suggesting that chromosome dynamics is dynamically arrested. The difference in density between the core and surface (Fig. 3E) suggests more architectural reorganization during XCI occurs on the surface, where the lower polymer density would allow for greater mobility ^59^. The lower density might also explain why most escape genes are located closer to the surface (Fig. 3C). In this case, the lower density would facilitate diffusion, mixing, and entry of necessary transcription factors into the chromosome territory. One method of investigating the change in global connectivity of the chromosome during inactivation is social network analysis, which examines coupling patterns. This analysis identifies connectivity hubs throughout the chromosome, enabling understanding of the change in interconnectedness (Fig. 4). We apply this analysis to the 3D Xa and Xi chromosome models, along with an unconstrained equilibrium polymer model, as a control. Here, connectivity of the chromosome is represented by a graph, where the degree of a node defines the number of contacts for that node and the local clustering coefficient measures the degree to which nodes cluster together. The degree centrality measures the number of connections a particular node has in the network. The results display a visible difference of information spreading on two types of models: unconstrained equilibrium polymers and Hi-C constrained chromosome models. The nodes in the former model are more uniformly connected with no obvious clustering, as opposed to Hi-C-constrained chromosome models, where we identify a number of nodes with high connectivity that form a single cluster, which may play an important role in information spreading.

**Fig 4.**
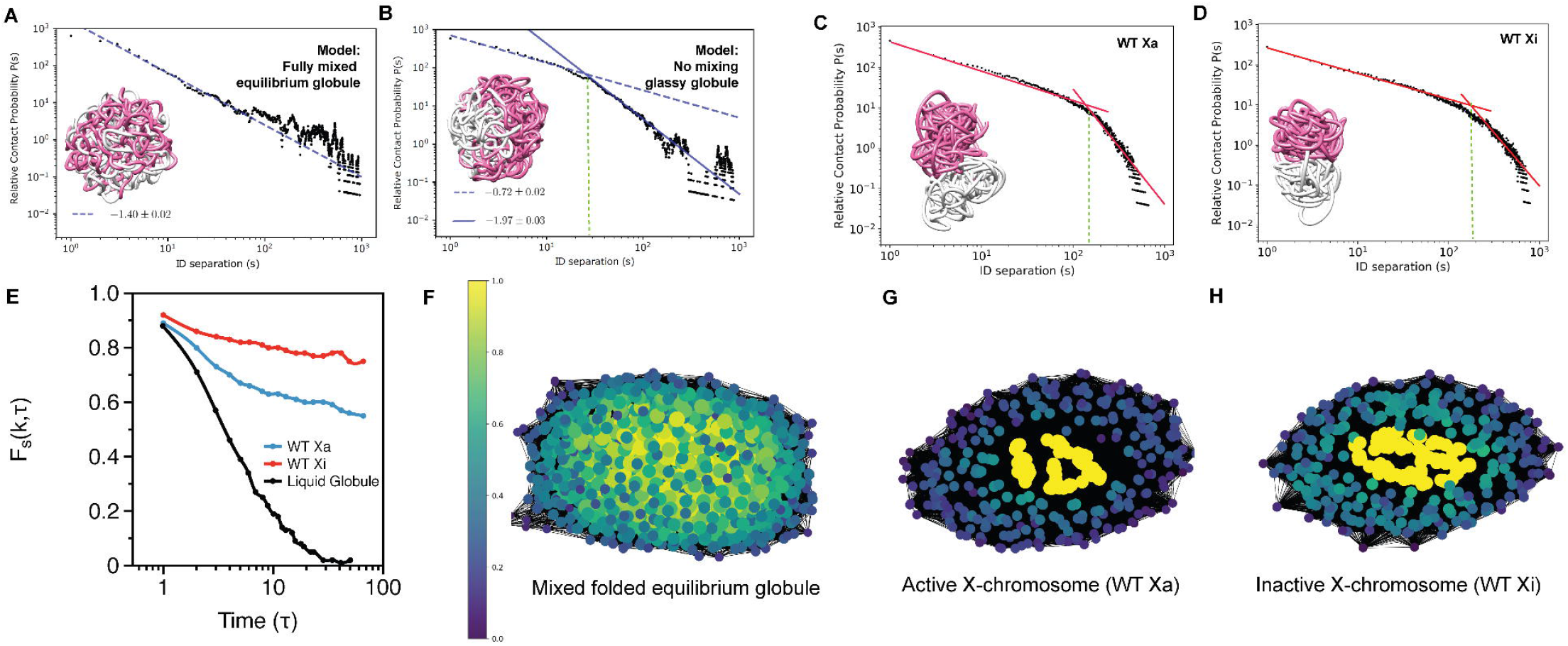
A link between mixing, compartmentalization and glassy dynamics. **(A)** A 3D model and contact probabilities for a fully mixed polymer chain. Pink and white and segments representing the centromeric and telomeric domains, respectively **(B)** A model of a dynamically arrested glassy polymer chain with no mixing, pink and white segments representing the centromeric and telomeric domains, respectively. For a fully mixed polymer in A, only one slope is present due to high degree of mixing, as opposed to two slopes in B, for a chain that does not mix, resulting in two modes of connectivity: short- and long range. **(C)** WT Xa model shows contact probabilities with two slopes, indicating that there is no mixing between the centromeric (pink) and the telomeric (white) domains and that there are two modes of connectivity: short- and long range. **(D)** WT Xi model shows contact probabilities with two slopes, indicating that there is no mixing between the centromeric (pink) and the telomeric (white) domains and that there are two modes of connectivity: short- and long range. **(E)** Self Intermediate Scattering Function F_s_(k, τ) demonstrates arrested dynamics behavior typical for a glass in both the active and the inactive X chromosomes. The F_s_(k, τ) decays to zero for a fully mixed equilibrium globule that does not have HiC constraints and therefore behaves as a liquid. **(F-H)** Profiles showing normalized betweenness centrality for **(F)** Folded equilibrium globule polymer chain **(G)** WT Xa chromosome model, and **(H)** WT Xi chromosome model. Values near 0 (purple) indicate low number of interacting neighbors. Values near 1 (yellow) indicate high number of interacting neighbors. Small node size indicates low number of interactions. Large node size indicates high number of interactions.

To quantify the level of mixing for the chromosome in a given state (as opposed to mixing during transitions between states discussed above) in detail, we examine the relative contact probability as a function of spatial separation. In principle, in a given state, a fully mixed chromosome would have a single slope (Fig. 4A), whereas a chromosome with no mixing would have a curve with two slopes and contain far fewer long-range interactions relative to the fully mixed configuration (Fig. 4B). Analysis of the Xa and Xi showed that both chromosomes have two slopes, indicating that relatively little mixing occurs in the Xa state (Fig. 4C) and that relatively little mixing occurs in the Xi state (Fig. 4D). Thus, both Xa and Xi polymers are each in a state of “dynamic arrest” rather than being freely mixing within their respective states. This is consistent with chromosome compartmentalization and loop formation as a result of HiC constraints, which allows for localized contacts within the A/B, S1/S2, or megadomain structures.

### Xist RNA spreading in 4D

Another powerful adaptation of our method is the ability to reconstruct the time-evolution of complex biological processes. We used our method to examine three dynamical processes. First, we analyzed the time evolution the X chromosome architecture during the Xa to Xi transition (Fig. 5). Interestingly, although the long-range interactions required for Xi megadomains are not present in the Xa Hi-C map, the regions corresponding to the proximal (white) and distal (purple) megadomains still undertook compact globular morphologies that were spatially separated from each other in 3D space. This separation persisted throughout XCI, occurring at days 0, 3 and 7 (Fig. 5A). To elucidate the mechanism of megadomain formation, we examined the connectivity within each megadomain. In the proximal megadomain at day 0, connectivity is relatively localized, with regions connected to each other that are nearby each other in sequence position. Coloring the chromosome by sequence position with red indicating regions nearer the centromeric end of the megadomain, blue indicating regions nearer the telomeric end, and white regions in between (Fig. 5B), red, white and blue are clearly separated at day 0. As the chromosome evolves to day 3 and day 7, progressively more mixing within the megadomain between red, white and blue occurs, indicating significantly more long-range interactions, consistent with the appearance of large ‘boxes’ of the megadomains on the empirically measured Hi-C map. The intra-megadomain mixing can be seen from two other 3D perspectives of the structure (Fig. 5C, S4). In the case of the proximal megadomain, similar intra-megadomain mixing is observed, with clear separation between red, white and blue at day 0 and progressively more mixing of red, white and blue at days 3 and 7 (Fig. 5D-E, S4). Thus, time evolution analyses of intradomain connectivity reveals a progression from mainly nearest neighbor local interactions typical of A/B compartments of the Xa to increasingly longer-range interactions that typify S1/S2 compartments and, ultimately, the megadomains of the Xi, all consistent with empirically derived Hi-C maps.

**Fig 5.**
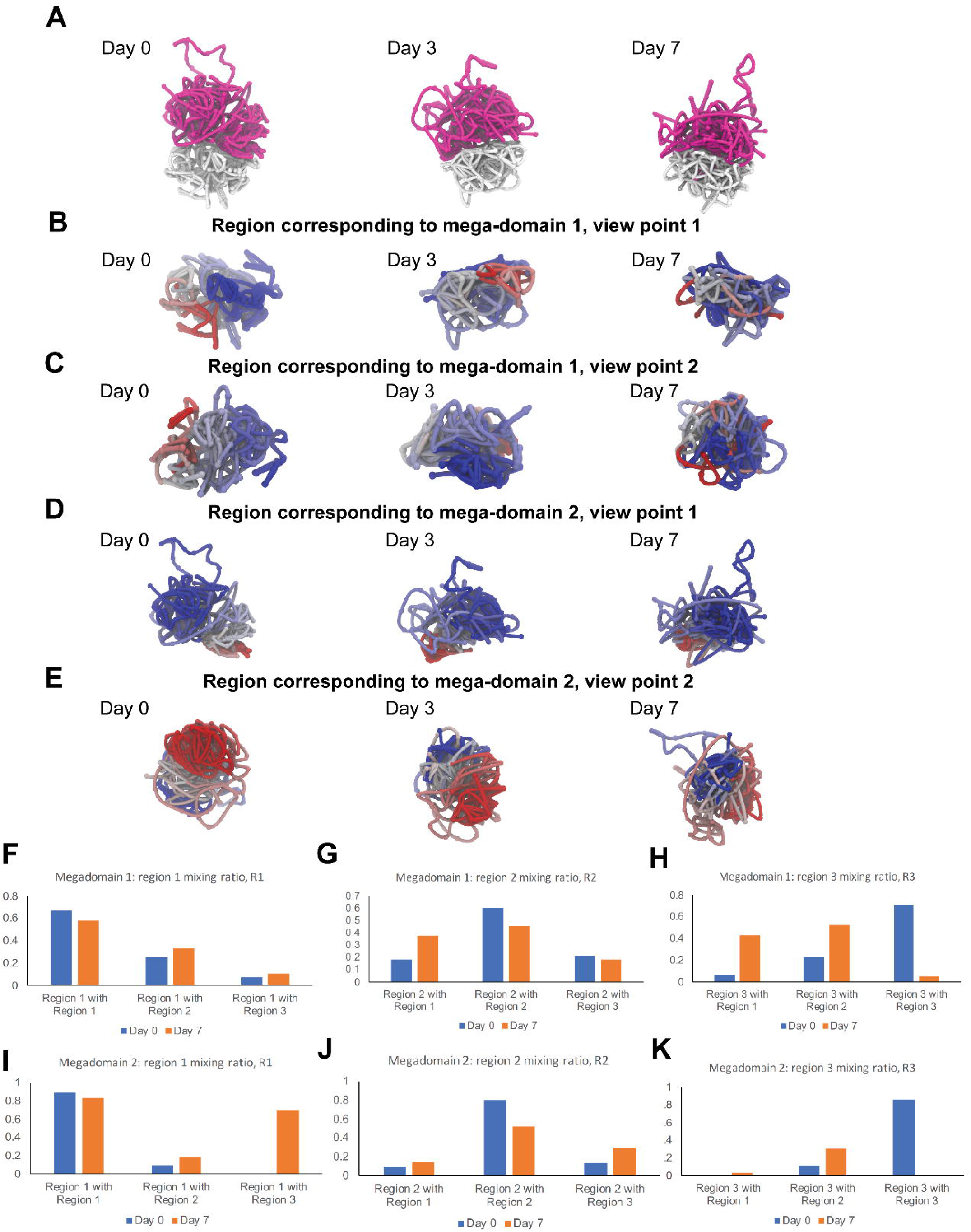
Intra-domain mixing of regions corresponding to the two megadomains of the X chromosome during X-inactivation. (A) 3-D reconstructions of the X chromosome based on Hi-C maps at day 0, day 3 and day 7 demonstrate that the two regions are separated in 3D physical space at days 0, 3, and 7. Pink, the region corresponding to megadomain 1 (positions 0 – 73 Mb); white, the region corresponding to megadomain 2 (positions 73 −166 Mb). (B) 3-D reconstructions based on Hi-C for region corresponding to megadomain 1 (positions 0 – 73 Mb), from same 3-D viewpoint as (A). This region undergoes significant mixing during X-inactivation. Chromosome is colored in a gradient from red to blue according to sequence position. At day 0, colors a separated. At day 7, colors are mixed, indicating intra-domain mixing. (C) Same as (B) but from a second viewpoint showing mixing at day 7. (D) 3-D reconstructions based on Hi-C for region corresponding to megadomain 2 (positions 73 −166 Mb), from same 3-D viewpoint as (A). This region undergoes significant mixing during X-inactivation. Chromosome is colored in a gradient from red to blue according to sequence position. At day 0, colors a separated. At day 7, colors are mixed, indicating intra-domain mixing. (E) Same as (D) but from a second viewpoint showing mixing at day 7. (F)-(K) Quantification of intra-domain mixing of regions corresponding to the two megadomains of the X chromosome during X-inactivation. (F) To quantify mixing within the region corresponding to megadomain 1, this region is divided evenly into three sub-regions (‘region 1’, ‘region 2’, and ‘region 3’). The mixing ratio for region 1, R1, is determined by computing the number of interactions between region 1 and another region divided by the total number of interactions. Self interactions (region 1 with region 1) decrease from day 0 to day 7. Intra-domain interactions (*e*.*g*., region 1 with region 2) increase from day 0 to day 7. (G) The mixing ratio for region 2, R2, is determined by computing the number of interactions between region 2 and another region divided by the total number of interactions. (H) The mixing ratio for region 3, R3, is determined by computing the number of interactions between region 3 and another region divided by the total number of interactions. (I)-(K) Same as (F)-(H) but for megadomain 2.

To quantify the intra-megadomain mixing of regions inside megadomains, we divide regions corresponding to megadomains evenly into three sub-regions (‘region 1’, ‘region 2’, and ‘region 3’ in Fig. 5F-K). The mixing ratio for each region, R_i_, is determined by computing the number of interactions within a cutoff radius, R_c_, between region i and another region divided by the total number of interactions. That is, for sub-region 1, three values of R_1_ are computed: one describing interactions of region 1 with itself, one describing interactions of region 1 with region 2, and one describing interactions of region 1 with region 3. In the case of megadomain 1 (Fig. 5F-H), as expected, self interactions (region 1 with region 1) decrease from day 0 to day 7. Intra-domain interactions (interactions of region 1 with region 2; interactions of region 1 with region 3) increase significantly from day 0 to day 7, demonstrating the occurrence of substantial mixing between these sub-regions within the megadomain during X-inactivation. Similar behavior is observed for R_2_ and R_3_, suggesting that all three sub-regions are significantly mixed, consistent with the graphical representations of the 3D reconstructions in Fig. 5. Likewise, we also observe substantial mixing between subregions of megadomain 2 (Fig. 5I-K). Taken together, our results demonstrate that, although the regions corresponding to the two megadomains appear spatially distinct in Xa and Xi, sub-regions within these megadomains which are spatially separated in Xa undergo significant mixing with each other during inactivation to arrive at the Xi configuration.

Second, we asked how Xist RNA – the noncoding transcript that induces whole chromosome silencing — spreads across the X-chromosome in relation to the large-scale changes of chromosome structure. Epigenomic methods such as “capture hybridization analysis of RNA targets (CHART)” have produced genome-wide binding maps for Xist RNA across time and yielded the hypothesis that Xist RNA spreads through ‘proximity’ transfer ^60,61^. However, in these previous studies, proximity referred to closeness in 1D sequence position along the polymer chain of the chromosome, rather than closeness in 3D physical space. Because 1D trends provide only part of the overall picture and 3D distances can be much shorter than 1D distances, it is imperative to examine proximal locations and spatial migration dynamics over a 3D Xi — a feat not yet modeled for Xist spreading. Here, by integrating Hi-C connectivity data with 1D CHART data, our 4DHiC pipeline now enables mapping the time-evolution of Xist spreading on a 3D model of the Xi.

We integrated data from a time-course Hi-C and CHART analysis ^34,51,60^ from the pre-XCI state (day 0) to mid-stage (day 4) and late-stage (day 7) XCI in wildtype cells (Fig. 6A) by superimposing the 1D signal onto a 3D structure of the chromosome. Our modeling indicated that the *Xist* gene localized at the surface of the active X-chromosome, specifically on the A compartment side of the chromosome (Fig. 6A, day 0, inset ii). Intriguingly, our modeling indicated that, although *Xist* resides in the A-compartment, it is located near the interface between the A- and B-compartments (Fig. 6A, day 4). Xist RNA was synthesized within the A-compartment and, at day 4, spread along the ipsilateral side to cover the proximal hemisphere of the Xi — the hemisphere corresponding to the A-compartment (Fig. 6A, red-colored polymer segments coinciding with the A-compartment in inset ii). The axis of the A/B compartments was perpendicular to the centromere-telomere axis (inset iii). A tomographic section through the Xi at day 4 revealed that Xist covered not just the surface of the A-hemisphere, but also regions within its core. By day 7, Xist had spread to the opposite hemisphere, now covering what were clearly separated A/B hemispheres of the former Xa. At the day 7 timepoint, however, it was also clear that the A/B compartments have ostensibly mixed together and condensed (Day 7, inset ii). Because Xist is known to spread with Polycomb repressive complex 2 (PRC2), we also examined the spreading dynamics of the Polycomb mark on trimethylated histone H3-lysine 27 (H3K27me3). The spreading of the H3K27me3 mark paralleled the migration of Xist RNA. Thus, Xist RNA, along with the epigenetic complexes that it recruits, emanates from a single point source in the A-compartment and diffuses in 3D space both through the chromosome and along its surface.

**Fig 6.**
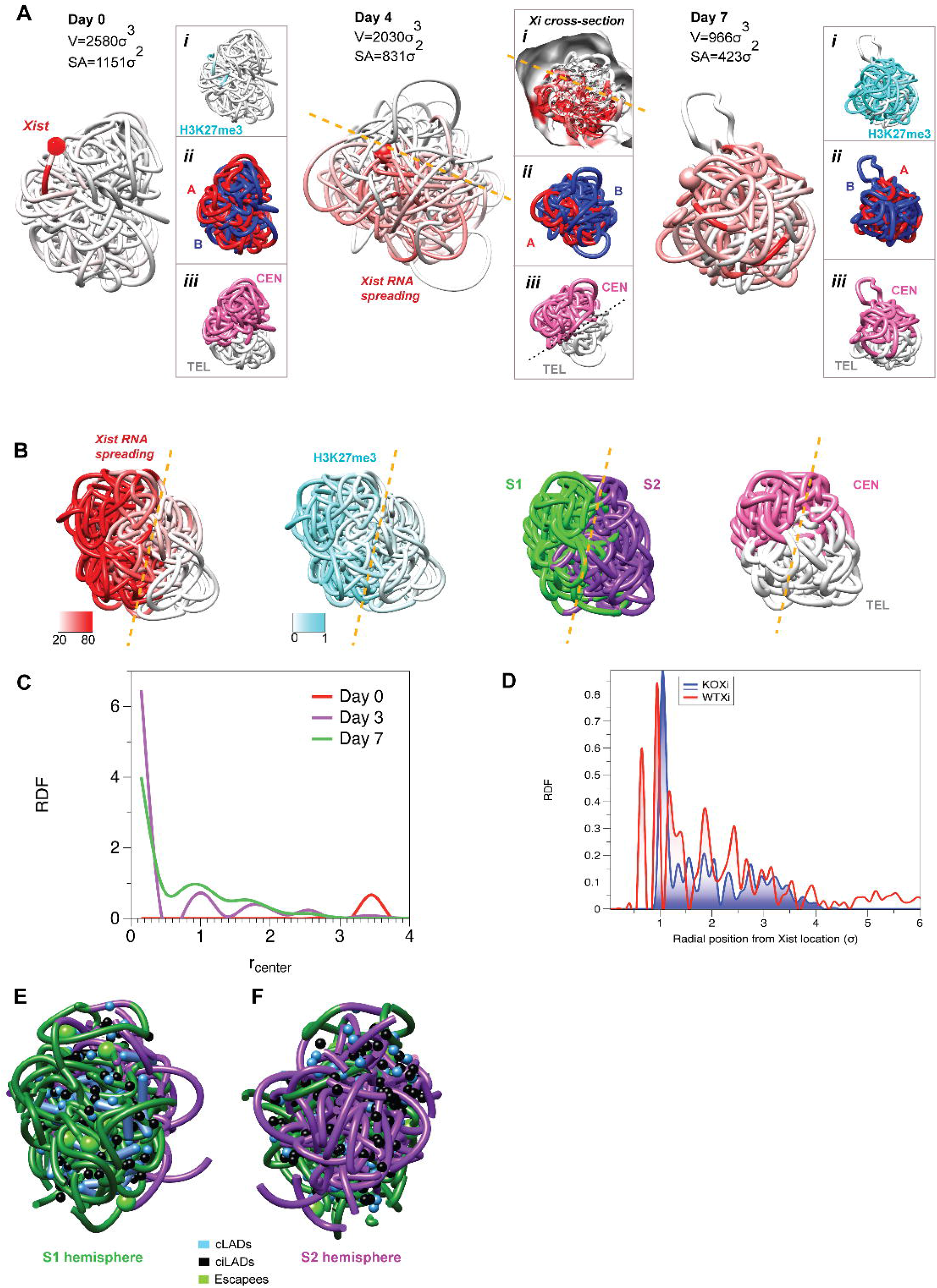
Xist RNA CHART - based RNA and methylation profiles in 3D in the WT and SMCHD1 knockout chromosomes. **(A)** 3D localization of Xist RNA (red) on the chromosome in pre-, early- and mid-XCI (day 0, day 3 and day 7, respectively). Insets: (i) localization of H3K27 methylation in the same orientation of the chromosome ii) localization of A and B compartments in the same orientation of the chromosome iii) localization of telomeric and centromeric domains in the same orientation of the chromosome. **(B)** Localization of Xist RNA, H3K27 methylation, S1/S2 compartments and centromeric vs telomeric domains in the SMCHD1 KO chromosome: both methylation and Xist occupy the S1 compartment that is orthogonal to the axis of centromeric and telomeric domains. **(C)** Radial distribution function as a function of radial position from Xist location for days 0, 3 and 7 (red, blue and green curves respectively)s. **(D)** Radial distribution function as a function of radial position from Xist location demonstrates the differences between Xist RNA spreading between the WTXi (red curve) and SMCHD1 knockout (blue curve) chromosomes. **(E)** Localization of constitutive lamin associated domains (cLADs), and constitutive inter-lamin associated domains ciLADs and escape genes on the S1 and **(F)** S2 hemispheres of the inactive chromosome in 3D as described in E.

Given the above, we next asked how the absence of SMCHD1 and failure to merge S1 and S2 compartments influenced the spreading dynamics of Xist RNA and H3K27me3. We mapped Xist CHART and H3K27me3 ChIP data onto the 3D reconstruction of the Xi in SMCHD1-deficient cells, and then projected the S1 (green) and S2 (purple) compartments onto the same model (Fig. 6B). Intriguingly, Xist RNA and H3K27me3 now only decorate one face of the Xi — specifically the S1 (green) compartment at day 7 (Fig. 6B) — at a time when Xist ordinarily would have already spread to the opposite face of the Xi (Fig. 6A). Thus, Xist stalls in the S1 compartment and fails to spread to the S2 half of the Xi. Likewise, H3K27me3 was observed to be enriched on the S1 side of the Xi, unable to continue onwards to the S2 side. While these data are consistent with previous 1D and 2D findings that Xist RNA and PRC2 become trapped in the S1 domain when SMCHD1 is lost ^51^, here we were able to visualize compartments in relation to the migration pattern of Xist RNA and PRC2 along the 3D surfaces of the A/B- and S1/S2-type of chromatin across time, thereby characterizing the process in 4D.

Here we were also able to study Xist RNA’s spreading through the radial distribution function (RDF). Whereas Xist spread outwardly from radial positions 0-6σ in WT cells, Xist could not do so in SMCHD1-deficient cells beyond 4σ (KS statistic=0.49, p=1.33e^-11^) (Fig. 6D). In the SMCHD1-deficient Xi, escapees partitioned into the S1 hemisphere, whereas constitutive lamin-associated domains (cLADs) and constitutive inter-lamin domains (ciLADs) were distributed across the S1 and S2 hemispheres (Fig. 6E-F).

Finally, we endeavored to encapsulate the above discoveries in a novel 4D simulation of time-dependent conformational changes in the X chromosome with simultaneous diffusion of Xist RNA particles around and through the chromosome. In particular, to visualize the formation of Xist RNA cloud around Xi during spreading and to investigate the spreading mechanism, we performed particle-based reaction diffusion simulations ^62^. This method is similar to the simulation method described above, but allows for additional processes, including chemical reactions, particle creation (*e*.*g*., transcription), and binding. Specifically, we simulated similar 3D transitions to those described above (from Xa to Xi) subject to the same Hi-C constraints, with the additional effects of RNA transcription, RNA diffusion and RNA binding to the chromosome. Consistent with our above findings, our reaction-diffusion model used the X-inactivation center (*Xic*) as the single point source from which Xist RNAs are transcribed (Fig. 7, purple sphere). As each bead in the chromosome represented 200 kb (consistent with the 200 kb Hi-C resolution), each Xist RNA (∼15 kb) was modeled by a single particle.

**Fig 7.**
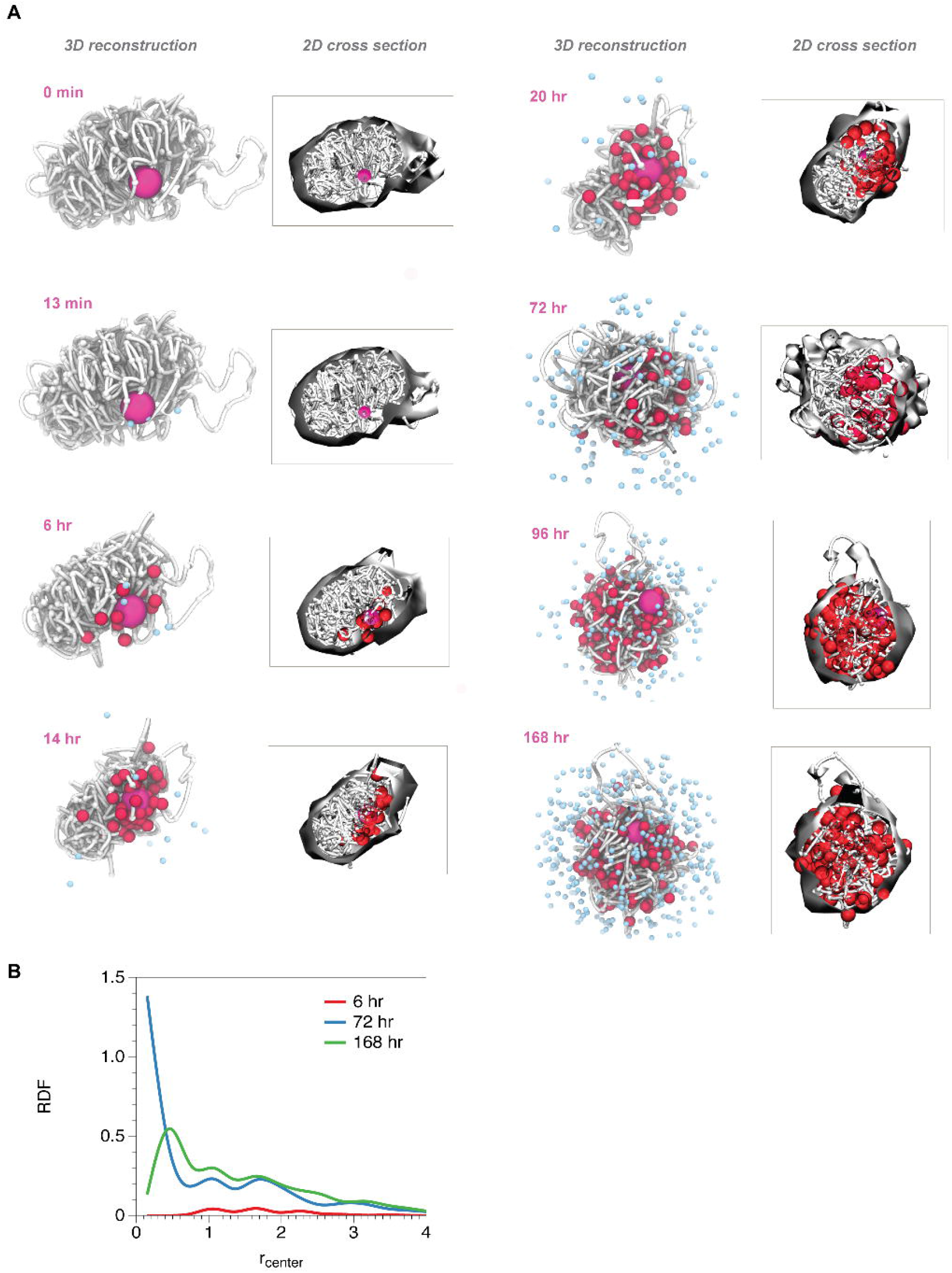
Time-evolution snapshots from particle-based reaction diffusion simulations demonstrate that Xist spreading can cover the X chromosome by diffusive processes alone. **(A)** Time evolution of chromosome structures and Xist RNA particles, with corresponding cross sections through the chromosome, displaying the interior of the chromosome (insets). White chain, X chromosome; magenta, X inactivation center (XIC); red, bound Xist RNA; light blue, diffusing Xist RNA. **At** time = 0 days, 13 minutes, newly transcribed Xist RNA particles visible (blue). At time = 0 days, 6 hours, Xist RNAs begin binding to experimentally determined Xist binding sites (red). At time = 0 days, 14 hours, RNA particles begin to penetrate interior of chromosome. At time = 0 days, 20 hours, RNA particles reach core of chromosome (inset). At **t**ime = 3 days, chromosome has undertaken large reorganization. At **t**ime = 4 days, 5 hours, RNA particles cover much of X chromosome. At **t**ime = 7 days, Xist RNA has covered majority of measured Xist binding sites, including those in the interior. **(B)** Radial distribution function (RDF) for bound Xist RNA particles for 6 hr, 72 hr, and 168 hr shows Xist RNAs gradually penetrate core of chromosome. X-axis is in units of σ.

Once transcribed, RNAs were allowed to freely diffuse in 3D space around and through the chromosome, subject to crowding by the cellular environment, emulated with spherical boundary conditions. RNAs then may bind back to the chromosome with a given rate, *k*_*binding*_, to designated Xist binding sites, as determined experimentally by our CHART experiments ^34,51^. In accordance with the above 3D reconstructions (Fig. 6), Xist RNA was simulated to initially diffuse from the proximal hemisphere, along the surface as well as through the internal core (Fig. 7, Movie 2). Simultaneously, the X chromosome undergoes large conformational changes during the transition from the Xa to Xi state (Fig. 7, Movie 2). Rates and parameters were chosen consistent with experimental observations, with the transition from Xa to Xi occurring on the time scale of 7 days and RNA spreading occurring over 7 days.

At very early times (0 days, 13 minutes), transcription events have begun. By 6 hours, 15 transcription events and 11 Xist RNA binding events have occurred. At 6 hours, the remaining four Xist RNA particles shown in the image have not bound and are still diffusing around the chromosome. By 14 hours (inset), RNAs have diffused into the intermediate layer of the chromosome (‘mantle’). At 20 hours (inset), significant penetration of the chromosome’s core has occurred. By day 3 (72 hr, inset), approximately half of the chromosome has been coated by Xist RNAs. At day 7 (168 hr), nearly all of the Xist binding sites on the chromosome are occupied by Xist RNAs. To quantify the Xist RNA penetration into the chromosome interior, we calculate the radial distribution function for Xist RNA particles bound to the chromosome, centered around the center of mass of the Xi chromosome, as a function of time (Fig. 7B). At 6 hr, the radial distribution function displays low density, consistent with the structure shown in Fig. 7A at 6 hr. By 3 days (72 hr), The RNA particles have covered about half of the surface of the chromosome and also penetrated about half of the chromosome’s interior, reaching its center, resulting in a large peak in the radial distribution function near the center of the chromosome (r_center_ = 0.25 σ), consistent with the structure shown in Fig. 7A at 72 hr. By day 7 (168 hr), the radial distribution peak shifts to r_center_ = 0.5 σ, consistent with a larger region of the core filling with RNA particles, as well as more RNAs binding towards the surface, as shown in Fig. 7A (168 hr). Taken together, the data suggest that crowding, and diffusion were sufficient to achieve rapid spreading of Xist over the full X chromosome. Specifically, as the *Xic* moved from its Xa to Xi position in 3D space, Xist RNAs quickly formed a cloud around the chromosome, maintained by crowding, and were positioned to cover the surface of the full chromosome (Fig. 7, Movie 2). Simultaneously, the RNA particles initiating at the *Xic* diffuse through the chromosome, coating the interior regions. A third effect occurs where RNA particles diffuse into solution around the chromosome and then enter the chromosome from surface regions distal from the *Xic*, allowing more rapid coating of the intermediate layer between the surface and core (‘mantle’). Control simulations that allow RNA diffusion but do not allow conformational change of the chromosome (*i*.*e*., ‘frozen’ chromosome) show less occupancy of Xist RNA binding sites on the chromosome, suggesting the specific conformational changes undergone by the X chromosome may help optimize the process of Xist coating through the various modes of Xist RNA binding. It is worth noting that the changes in 3D conformation of the X chromosome do not occur independently of spreading. We anticipate that 3D changes are highly interconnected with Xist and Polycomb spreading, with the latter also helping to propagate the former. As more experimental evidence elucidates this relationship, we hope to incorporate such interdependency in our models.

## Discussion

By combining 2D Hi-C experiments with polymer physics models and reaction-diffusion kinetics, we have constructed 4D models to demonstrate the organization and the dynamical properties of the X chromosome during the transition from the active to the inactive state. To validate each 3D reconstruction of the chromosome, we compute simulated 2D Hi-C maps. We find the simulated maps are highly correlated with the empirically measured Hi-C maps. We note that each empirically measured Hi-C map represents an average over cells, giving interactions and patterns of interactions that are most prominent over thousands of cells. Likewise, each of our 3D reconstructions is determined directly from a measured Hi-C map and is thus an abstract object depicting an average architecture that is ‘conserved’ over cells. From a physics standpoint, for any given set of interaction constraints, there may be a quite large ensemble of polymer configurations consistent with the constraints, especially for a non-equilibrium system such as a chromosome driven by the events occurring during the cell cycle. However, due to the high number of interactions caused by relatively dense packing into the nucleus (*i*.*e*., the connectivity matrix is densely populated), the ensemble is relatively small, suggesting that the 3D reconstructions depict an accurate reflection of the chromosome’s architecture in the cell, making such models valuable for studying XCI and chromosome dynamics in general.

From our 3D modeling, we have learned details that could not be deduced from 2D datasets. For the Xi, several layers of information have been concealed, and new methodologies have been needed to unveil these layers without disrupting chromosome folding. Previous studies had suggested that the original underlying structures of the Xa are destroyed when the chromosome is reorganized to the inactive state. However, by being able to identify and visualize specific 3D relationships and by studying cross sections through the chromosome, our models, based directly on the empirical Hi-C data, demonstrate that neither the A/B compartments nor the S1/S2 structures are wholly destroyed within the Xi superstructure. In fact, they appear to be merely refolded like origami and therefore hidden underneath the viewable surface of the Xi. This is critical information from the perspective of chromosome dynamics, as indeed it supports the architecturally more favorable model of large-scale structures being progressively built upon, rather than being unraveled and rebuilt during XCI. Previously, because of the limitation of viewing 2D information alone, the Xi gave the appearance of being unstructured, apart from its two megadomains. However, our new data, taken together with published studies, underscores the fact that the Xi is not unstructured, but rather contains structural remnants from earlier in the chromosome’s life cycle.

Our study provides cues on chromosome structure, allowing us to identify three layers of chromosome dynamics – slow dynamics in the dense core, intermediate layer (‘mantle’), and fast dynamics on the surface of the chromosome. Intuitively, this is coupled to contact density, implying that a folded chromosome is analogous to a glassy polymer and the role of surface effects is highly relevant to chromosome function, including interchromosomal interactions.

From our 4D study (*i*.*e*. the time series of 3D structures), we have also learned that the *Xist* gene — though located in the A-compartment — resides near the A/B interface. During XCI, Xist RNA originates in the A-compartment, and spreads through both the surface and the core of the A-compartment initially. Xist RNA then depends on SMCHD1 to migrate across the A/B interface to spread over the B-compartment. Within both hemispheres, Xist binds both surface and core chromatin. Our 3D analysis provides an effective way of tracking the spreading of Xist RNA, showing that the spreading resembles a diffusive wave-front emanating from a point source, the Xist locus. In the SMCHD1 knockout case, the wave-like propagation halts mid-way, effectively ‘polarizing’ the chromosome, only half of which gets covered by Xist RNA.

Notably, from 2D contact maps alone, Xist’s coverage of chromosome surface versus its core could not be determined. It is also notable that escapees reside near or at the surface of the Xi where the chromosome is less compact and where greater mobility is possible. Our deductions (Fig. 6, 7) are consistent with the wealth of published 2D contact maps ^8,34-36,38,51,63,64^ and microscopy data that have yielded much valuable information about locations of select genes within the Xi territory ^39-45^. The picture emerges of a well-organized but dynamic Xi during its formation and establishment. Taken together, our results are consistent with a gradual accumulation of structural elements throughout the process of X-inactivation, whose 4D architecture (*i*.*e*., the temporal series of 3D chromosome configurations) may aid in accelerating Xist diffusion and binding throughout the chromosome, ultimately resulting in rapid Xist coating and shutdown of the X-chromosome. Our 4DHiC pipeline is transferable to any other Hi-C dataset and can be used to model the organization of an entire nucleus.

## Methods

### 4DHiC Pipeline

Simulations described in this work were performed using the Large-Scale Atomic/Molecular Massively Parallel Simulator (LAMMPS)^65^ and Reaction Diffusion Dynamics (ReaDDy2)^66^. In the case of the LAMMPS simulations, the initial structure consisted of N=833 beads corresponding to ∼167 Mb. The initial configuration forms an open coil chain as a result of a self-avoiding random walk (SARW). Experimental Hi-C constraints were used directly such that connected pairs (virtual HiC bonds) were required to form harmonic bonds between interacting particles, and those were directed to form connected pairs prior to production simulations. Once all the bonding constraints were satisfied, the convergence was measured by comparing the experimental contact maps to the simulated ones using the Pearson correlation coefficient. To help verify the simulated contact maps from our 3D models, our MultiSim approach (described below) was used (Fig. S5). The bonds were preserved during production simulation steps and the structure was subjected to Brownian dynamics with implicit solvent. Defined for the simulation were pair interactions between bonded particles using FENE and Lennard-Jones potentials:

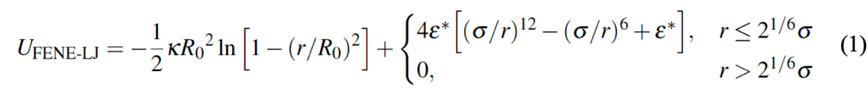

where σ is a dimensionless quantity that characterizes distance, and the optimal parameter set of the maximum bond length *R*_0_ = 15σ and the spring constant *κ* = 30*ε*^*^/σ^2^. We choose the repulsive LJ strength *ε*^*^ = 1 in non-dimensional units for this bonded potential, which makes the equilibrium bond length *r*_bond_ = 0.99σ yet allowing the bond to be stretched up to 15σ. This deviation from the classical Kremer-Grest Model is important for the functionality of the presented workflow. Allowing for large extensions is what allows one to generate models with unmappable regions where atoms have no contacts and therefore stay ‘looped out’, requiring large extensions since the rest of the chromosome rapidly compactifies. This modification has no effect on the actual structure because all consecutive particles are found within 1σ from each other, and their long-range connectivity is dictated by the virtual HiC constraints.

For nonbonded atoms, only the repulsive part of the Lennard-Jones interaction potential was used:

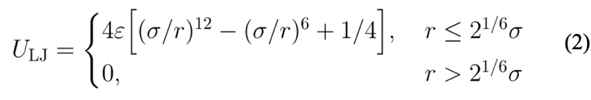

Finally, we used harmonic constraints originating from experimental Hi-C data, *U*_Hi−C_ = K(*r* −*r*_0_)^2^, where K=1 *ε/* σ^*2*^ and r_0_=2.2σ. This ensures that the initial random structure of the polymer chain converges and satisfies the constraints originating from Hi-C experiments. As a result of parameter studies, r_0_ = 2.2σ was found to be the optimal value such that a large number of connected particles may be accommodated without overlapping particles which would be unfeasible for the simulation. The average simulation temperature was controlled by the Langevin thermostat (kept constant at T_start_ = T_end_ = 1 in reduced units, with the damping coefficient set to 1 τ^-1^. A timestep of 0.01τ was used, where τ is the reduced (Lennard-Jones) time-a measure of how long it takes for the particle to move across its own size, defined as 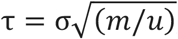, where *m* is the characteristic mass, and *u* is the intrinsic energy of the system that is the same as parameter *ε\** in the spring constant κ. Since the total number of particles within a sphere *r*_min_ includes the particle itself, we define the average number of contacts in the total system, for a given contact radius, as *z = (Ω*−*N)/N*, where *Ω* is the total number of contacts in the whole system for a given contact radius.

### Assessment of Chain Connectivity

The algorithm for generating a contact map from the polymer simulation is straight forward: for each consecutive monomer, we list all other monomers that are within the chosen contact radius. Obviously, the choice of this radius is somewhat arbitrary, and for simulation-based contact maps we regarded monomer *m*_*i*_ and monomer *m*_*j*_ as being in contact when they were found within the radius of 3σ, which was found to be the best to reflect the similarity between experimental and simulated contact matrices. The *off-diagonal* contacts are symmetric about the main diagonal, revealing the essential features of the collapsing polymer. Experimental Hi-C data contains unmappable regions where DNA sequence is repetitive, and it is impossible to know the exact connectivity of certain segments. In our model, the unmappable regions ‘loop out’ from the final folded state due to the fact that these regions have no specified Hi-C constraints. These regions are excluded from any further analyses.

We investigated the effect of the finite size of a globule by considering how the number of contacts and displacement of particles varies as a function of radial distance from the center of mass *r*_center_. The volume occupied by a collapsed globule can be divided into concentric spherical shells of thickness *d*_r_, centered on the center of mass. Each particle can be placed into a shell based on the position of its center. For every particle, the number of contacts is calculated, as well as the displacement from its position at the chosen reference time. The average number of contacts and the RMS displacement is calculated across all particles within each shell. It is then instructive to consider the radial profile by plotting these quantities against the average radius of each corresponding shell. A suitable number of shells is chosen such that there is a sufficient number of particles in each shell to calculate reliable averages; eight shells were used for a chain of N=833. Using spherical shells is naturally most accurate for spherical globules where there is a relatively uniform angular distribution of particles for a given radius; significantly non-spherical globules could introduce anomalies. To compute accessible surface areas, we used the Lee-Richards rolling ball algorithm for a ball of radius of 1.4σ. The rolling ball algorithm is a method that creates a mesh of points equidistant from each monomer of the chromosome and uses the number of these points that are solvent accessible to determine the accessible surface area. The points are constructed at a radius beyond the monomer radius (1.4σ), which is effectively similar to ‘rolling a ball’ along the surface. All points are checked against the surface of neighboring monomers to determine whether they are accessible. The number of points accessible is multiplied by the portion of surface area each point represents to calculate the Accessible Surface Area.

### Contact Probability Scaling

Contact probability scaling can only be straightforwardly applied to systems representable as linear polymer chains, where each segment’s position in the chain sequence is known. The contact probability, *P*(*s*), is defined for every possible arc-length separation, *s*, along the chain. It is given by the ratio of the number of contacts, *N*_*c*_, whose contacting particles have separation *s*, to the total number of possible contacts *N*_*p*_ For an equilibrium globule, *N*_*p*_ is proportional to a quantity *s*^1/2^, at the same *s*:

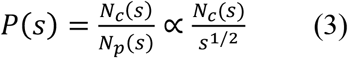

Structures with similar fractal dimensions but different conformations and topologies can then be distinguished by differences in the gradients obtained from a plot of log(*P*(*s*)) against log(*s*). Note that the value of the constant of proportionality in the definition of *P*(*s*) is irrelevant since we are only interested in gradients of this curve. Our results demonstrate that contact probability scaling can provide insights into types of connectivity such as local (unmixed globular regions) or global structures (fully mixed chains).

### MultiSim Approach

In our MultiSim approach (Fig. S5), a number of structures with constraints chosen at random and equally distributed across 10 independent simulations were averaged. In contrast to the 4DHiC approach, this method satisfies a larger number of constraints due to the fact that there is a smaller number of bonds per generated structure, allowing for increased conformational freedom. Each MultiSim simulation has the full-length chromosome (N=833), but each simulation only contains 1/10 of total bonds, randomly distributed across simulations without repetitive occurences. 4D HiC simulation alone produces tractable results that were used to draw general conclusions about the different states of the X-chromosome. It has to be emphasized that the experimental Hi-C data is an average of hundreds of thousands of cells and does not as such represent a single structure that can be identified experimentally. To address this issue, we set up simulations using a randomly chosen subset of empirically determined Hi-C constraints, dividing them equally into 10 simulations, after which all the data from separate simulations was combined in order to reconstruct the 3D state. This way, while studying the individual simulations does not yield all the necessary information that could be used for data interpretation, the combined results are nearly identical to both experimental and ensemble averaged data, with R_Pearson_=0.91, increasing the target virtual HiC bonding by 10% in comparison to 4DHiC. To increase the specificity of virtual simulated bonds in reconstructing a 3D structure with the empirical HiC contacts using MultiSim, we incorporated contact frequency as another factor influencing the polymer chain condensation. Chromosome conformation capture techniques can capture the relative occurrence of specific contacts within the chromosome sequence. In analyzing this data, we follow with the logic that contacts which appear more frequently within the HiC data are a result of high affinity regions that are more critical to the overall condensed structure of the X-chromosome as compared to regions with lower contact frequencies, which may be the result of random chance. Since there are many possible 3D solutions to any given contact map, it is important to prioritize the formation of contacts which appear across a vast majority of empirical chromosome structures. We adjusted the model by simulating higher frequency contacts with stronger bonds, implementing a system of binning frequency ranges with a chosen bond energy in simulation. Two models were tested in determining whether introducing frequency would increase contact accuracy. In the first model, contact frequency ranges obtained from HiC data were sorted into 11 equally sized bins, each bin representing a different bond strength in simulation. Here, we used linearly increasing bond strengths per bin to correlate with each frequency range. In the second model, we instead used exponentially increasing bond strengths per bin. Based on these simulations, there was no significant difference in contact accuracy as a result of introducing linearly increasing or exponentially increasing bond strength binning.

#### Social Network Analysis of Contact Matrices

Social network analysis used in this work was based on symmetric networks, using the symmetry of chain connectivity in contact matrices (*i*.*e*., if monomer 1 is connected to monomer 25, the reverse is also true). The degree of a node defines the number of connections a node has. The local clustering coefficient is a measure of the degree to which nodes in a graph tend to cluster together. The degree centrality is a measure of the number of connections a particular node has in the network. It emphasizes the fact that ‘crucial’ nodes have many connections. This analysis was performed using NetworkX v2.0 and Python 3.5.

### Particle-based Reaction Diffusion simulations of RNA spreading around the X chromosome

The ReaDDy2 particle-based reaction diffusion simulation package was used to simulate RNA spreading around and through the X chromosome ^62^. To perform particle-based reaction-diffusion simulations, the Langevin equation is solved,

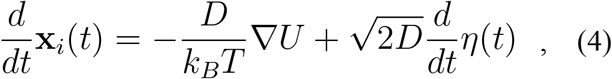

using the integrator,

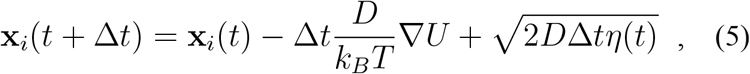

where **x**_*i*_ is the position of particle *i, t* is the time, *U* is the pairwise potential energy function for interactions, *k*_B_ is the Boltzmann constant, *T* is the temperature, *η(t)* is the noise function and D is the diffusion coefficient. In addition, equations governing reaction kinetics are used to simulate the creation of new particles as well as binding interactions. Transcription of Xist RNA can be described with the unimolecular reaction,

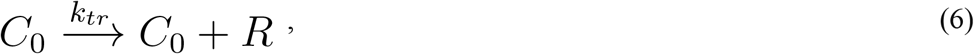

with differential rate law,

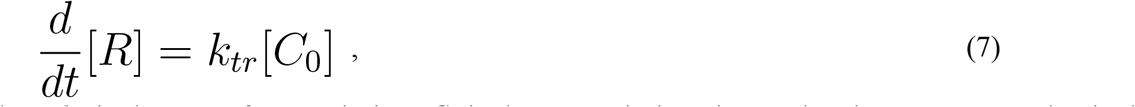

where *k*_*tr*_ is the rate of transcription, *C*_*0*_ is the transcription site on the chromosome, and *R* is the transcribed Xist RNA. In practice, a new Xist particle is created with rate *k*_*tr*_. *Xist* RNA binding to the chromosome is described by the bimolecular reaction,

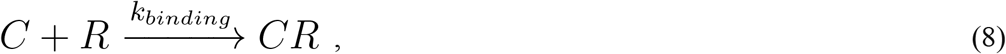

with differential rate law,

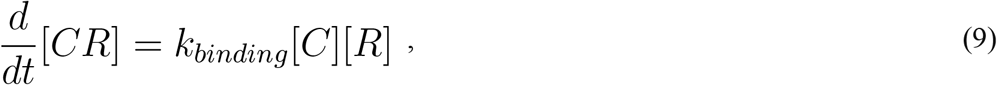

where *k*_*binding*_ is the rate of binding, *C* is the Xist binding site on the chromosome, *R* is the Xist RNA and *CR* is the complex of RNA bound to the chromosome. In practice, RNA binding is simulated with molecular microreactions between the Xist RNA particle *i* and the chromosome monomer particle *j*. Here, when the educts (RNA particle and chromosome mononmer particle) are located inside a cutoff radius of each other, they form an encounter complex with reaction rate, *k*_*binding*_. Specifically, if the RNA particle *i* enters an encounter radius *r*_*ij*_ of chromosome monomer particle *j*, then a binding reaction with rate *k*_*binding*_ occurs with Poisson probability,

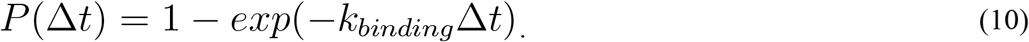

While this is a simplified model, it allows us to explore, for the first time, the process of RNA transcription and RNA binding to the chromosome during conformational changes of the chromosome for the first time. The basic parameters of the model are the radius of chromosome monomers, *r*_*C*_, the radius of the RNA particle, *r*_*R*_, the rate of binding of the RNA, *k*_*binding*_ and the rate of production of the RNA, *k*_*tr*_. Experimentally, none of these quantities is known with high precision for the specific system considered. While we use radius *r*_*C*_ corresponding to the resolution of the HiC map (200 kb), it is difficult to approximate the volume occupied by 200 kb of chromatin since each 200 kb segment of chromatin could occupy different amounts of volume due to differences in conformation as well as nucleosome and protein occupancy. Likewise, in the case of *r*_*R*_, there is no information available regarding the volume occupied by Xist since little is known about its 3D structure. Even less is known about the exact binding rate of Xist to the chromosome. While many estimates of an average *k*_*tr*_ exist, the precise value of *k*_*tr*_ for Xist in the context of mouse neural progenitor cells (NPCs). during X-inactivation is not known. Given the uncertainty in experimental data, we use a simple approximation of *r*_*R*_ ∼ *r*_*C*_. We adjusted the rates such that Xist spreading across the chromosome occurs in the experimentally determined time of 7 days. To emulate crowding by the cellular environment surrounding the chromosome, we employ spherical boundary conditions.

To simulate the conformational change of the X chromosome from Xa to Xi, we use a method based on steered molecular dynamics ^67 68-70^. The initial (active X) and target (inactive X) were derived from the Hi-C based structures discussed earlier. Steered molecular dynamics, which was originally used to model atomic force microscopy experiments, achieves conformational changes by coupling to an external force, often implemented as a harmonic potential. We note that the concern that thermodynamic potentials cannot, even in principle, be obtained from irreversible processes was disproved by Jarzynski, who proved the identity ^71,72^, <*exp*[- *W*/k_B_*T*]> = *exp*[− *ΔF*/k_B_*T*], which connects the ensemble average of an exponential of the total work *W* performed on a system during the non-equilibrium transition from one state to another to the free energy difference Δ*F* between the two states. In the limit of exhaustive sampling, steered molecular dynamics is, in principle, capable of producing equilibrium free energies of conformational changes ^73^. An advantage of this method is that it ensures that active and inactive states are consistent with the experimentally determined Hi-C data.

Raw next-gen sequencing data and processed files for Hi-C, ChIP-seq and CHART-seq have been deposited in Gene Expression Omnibus (https://www.ncbi.nlm.nih.gov/geo/) under accession number GEO: GSE99991.

## FIGURE CAPTIONS

**Movie 1**. 3D reconstruction of the Wild Type (WT) Xa chromosome model. Pink— centromeric domain; white— telomeric domain.

**Movie 2**. Time-evolution from particle-based reaction diffusion simulations demonstrate that Xist spreading can cover the X chromosome by diffusive processes alone. White chain, X chromosome; magenta, X inactivation center (XIC); red, bound Xist RNA; light blue, diffusing Xist RNA. See figure 5 for details.

## Acknowledgements

We thank members of the Sanbonmatsu and Lee Labs for stimulating discussions. This work was funded by LANL LDRD 20210082DR and LANL LDRD 20210134ER to K.Y.S and A.L., F31HD100109-01 to A.J.K., and the National Institutes of Health (R01-HD097665) to J.T.L. We are grateful to LANL Institutional Computing for their generous support.

## Author Contributions

Conceptualization: Anna Lappala, Jeannie T Lee, Karissa Y. Sanbonmatsu

Formal Analysis & Data Interpretation: Anna Lappala, Kevin Tan, Hunter Michalk, Karissa Y.

Sanbonmatsu, Jeannie T. Lee

Funding acquisition: Karissa Y. Sanbonmatsu, Jeannie T. Lee

Methodology: Anna Lappala, Karissa Y. Sanbonmatsu

Supervision: Jeannie T. Lee, Karissa Y. Sanbonmatsu

Writing – original draft: Anna Lappala

Writing – review & editing: Jeannie T. Lee, Anna Lappala, Karissa Y. Sanbonmatsu, Andrea Kriz, Chen-Yu Wang

## Competing Interests

Authors declare no competing interests.

## Data and materials availability

All data and code used in the analysis are available upon reasonable request.

## Supplementary Materials

Materials and Methods

Figures S1-S5

Tables S1

Data

Analysis scripts

## Notes

### Competing Interest Statement

The authors have declared no competing interest.

